# Transcriptomics of independent CRISPR-edited cell lines reveal ciliary-specific ARL13B dependent changes

**DOI:** 10.64898/2026.08.13.744725

**Authors:** Olivia Morrison, Tamara Caspary

**Affiliations:** Genetics and Molecular Biology Graduate Program, 615 Michael St., Suite 301 Atlanta, GA, 30322; Department of Human Genetics 615 Michael St., Suite 301 Atlanta, GA, 30322

**Keywords:** ARL13B, Primary cilia, Ciliary signaling, Kidney epithelial cells, Renal cystogenesis, RNA sequencing, Ciliopathy

## Abstract

Primary cilia coordinate signaling pathways that regulate tissue homeostasis and development, and defects in cilia contribute to numerous ciliopathies. However, the transcriptional consequences of disrupting ciliary protein localization remain poorly defined. ARL13B is a cilia-enriched regulatory GTPase required for ciliary trafficking and signaling. The ARL13B^V358A^ variant is undetectable in cilia yet retains known biochemical functions, providing a unique model to investigate the functions of ciliary ARL13B independently of ciliogenesis. To define transcriptional programs associated with loss of ciliary ARL13B, we generated two independent *Arl13b^V358A/V358A^* kidney epithelial cell lines and matched rescue lines. The ARL13B^V358A^ mutation did not affect ciliation frequency or cilia length but altered ciliary protein composition, including loss of ARL3 and INPP5E localization and increased accumulation of TULP3. RNA sequencing revealed expression changes in genes associated with ciliary biology, mechanotransduction, epithelial organization, and kidney-related phenotypes. Despite similar ciliary phenotypes, the independently-derived, mutant clones displayed substantial transcriptomic heterogeneity, highlighting a potential source of variation in CRISPR-based transcriptional studies. By integrating data from the independent mutant and rescue clones, we identified a high-confidence set of 131 genes whose expression reproducibly tracked with loss and restoration of ciliary ARL13B. Together, these findings demonstrate that ciliary ARL13B is required to maintain normal ciliary composition and gene expression programs and underscores the value of multi-clone, rescue-based experimental designs for robust transcriptomic analyses.

**Summary for Reviewers:** This study examined how excluding the protein ARL13B from primary cilia affects kidney epithelial cells. The researchers created two independent cell lines carrying a modified form of ARL13B, along with matched rescue cell lines. The findings show that ciliary ARL13B helps maintain normal ciliary composition. By comparing the cell lines, the researchers identified a high-confidence set of genes associated with loss of ciliary ARL13B. By highlighting the importance of using independent gene-edited clones and rescue-based controls, these results advance understanding of how cilia regulate kidney cell function and provide guidance for designing robust transcriptomic analyses.

## Introduction

Primary cilia are specialized sensory organelles that integrate diverse signaling pathways governing development, tissue homeostasis, and disease (Bangs and Anderson 2017; Hilgendorf et al. 2024). These solitary, microtubule-based projections extend from the cell surface and originate from a modified centriole, the basal body (Sorokin 1968). The ciliary axoneme is composed of microtubule doublets surrounded by a specialized membrane that is continuous with, yet compositionally distinct from, the plasma membrane (Garcia-Gonzalo et al. 2015; Chavez et al. 2015). Primary cilia concentrate ion channels, G protein-coupled receptors (GPCRs), and signaling effectors within a confined membrane-bound compartment (Nachury and Mick 2019). This organization increases the local concentration of signaling molecules, enabling sensitive detection and efficient amplification of extracellular cues (Ott and Lippincott-Schwartz 2025). In addition to promoting efficient signal detection, this compartmentalization allows cells to spatially restrict signaling activities from those occurring in the cytoplasm, thereby fine-tuning downstream cellular responses (Truong et al. 2021; Mukherjee et al. 2016).

The signaling functions of primary cilia depend on regulated mechanisms that control protein entry, exit, and retention within the organelle. At the ciliary base, the transition zone serves as a selective diffusion barrier (Garcia-Gonzalo et al. 2011; Chih et al. 2011), while intraflagellar transport (IFT) and other ciliary machinery mediate the trafficking of proteins into and out of the ciliary compartment (Ou et al. 2005; Kozminski et al. 1993; Jin et al. 2010; Nachury et al. 2007). Together, these systems establish and maintain a specialized signaling environment that supports pathways essential for tissue development and homeostasis, including vertebrate Hedgehog (Hh), Wnt, cyclic AMP, and mechanosensory signaling (Huangfu et al. 2003; Nauli et al. 2003; Wallingford and Mitchell 2011; Siljee et al. 2018; Hilgendorf et al. 2016). Reflecting the broad physiological roles of primary cilia, disruption of ciliary structure or signaling give rise to a diverse group of multisystem disorders known as ciliopathies (Reiter and Leroux 2017; Hilgendorf et al. 2024). These disorders affect numerous organ systems, including the kidney, retina, liver, skeleton, and central nervous system, and encompass diseases such as polycystic kidney disease, Bardet-Biedl syndrome, Joubert syndrome, and Meckel syndrome (Reiter and Leroux 2017; Hilgendorf et al. 2024).

Although the role of primary cilia as signaling organelles is well established, defining how changes in the abundance and localization of specific ciliary proteins influence downstream transcriptional programs remains a significant challenge. One major limitation is that many ciliary proteins perform essential structural roles in ciliogenesis and ciliary maintenance (Pazour et al. 2000; Kozminski et al. 1993; Murcia et al. 2000). Consequently, loss of these proteins often disrupts the entire organelle, making it difficult to distinguish transcriptional effects arising from the absence of specific ciliary signaling activity from those resulting from complete ciliary dysfunction. Experimental systems that selectively alter the subcellular localization of ciliary proteins while preserving their biochemical function offer a powerful strategy to overcome this challenge (Zuo et al. 2019; Corbit et al. 2005). Such approaches make it possible to determine how the localization of specific ciliary proteins influences cellular transcriptional programs while minimizing the confounding effects of complete ciliary loss.

ARL13B is a regulatory GTPase that is highly enriched in the ciliary membrane, where it regulates protein trafficking, phosphoinositide composition, and the ciliary enrichment of proteins such as ARL3 and INPP5E (Humbert et al. 2012; Larkins et al. 2011; Sun et al. 2004; Gigante et al. 2020; Fujisawa et al. 2021; Qiu et al. 2021). In addition to binding and hydrolyzing GTP, ARL13B functions as a guanine nucleotide exchange factor (GEF) for ARL3 (Gotthardt et al. 2015). Cells lacking ARL13B exhibit increased ciliary accumulation of the phosphoinositide-binding trafficking adaptor TULP3, consistent with disruption of ARL13B-dependent control of ciliary phosphoinositide composition and membrane protein trafficking (Garcia-Gonzalo et al. 2015; Gigante et al. 2020; Mukhopadhyay et al. 2010; Nozaki et al. 2017). Loss of ARL13B leads to abnormal cilia traffic of vertebrate Hh pathway components and Hh transcriptional response (Caspary et al. 2007; Larkins et al. 2011). Loss of ARL13B, as occurs in the *Arl13b^hennin(hnn)^* mouse, also disrupts ciliogenesis and alters ciliary architecture (Caspary et al. 2007).

Ciliary localization of ARL13B is required for normal renal homeostasis (Van Sciver et al. 2023). The ARL13B^V358A^ (ARL13B^A^) variant retains its known biochemical activities but is undetectable in cilia, thereby providing a tool with which to isolate ciliary ARL13B function (Gigante et al. 2020; Mariani et al. 2016). *Arl13b^V358A/V358A^* (hereafter called *Arl13b^A/A^)* mouse embryonic fibroblasts display abnormal cilia, disrupted ciliary localization of ARL3 and INPP5E, but normal vertebrate Hh pathway response (Gigante et al. 2020). This demonstrates that ciliary and cellular ARL13B possess distinct functions. *Arl13b^A/A^* mice develop cystic kidneys, but the transcriptional response to this perturbation has not been characterized (Van Sciver et al. 2023). Defining the transcriptional consequences of specifically excluding ARL13B from cilia is important for understanding how ciliary localization of ARL13B contributes to epithelial cell behavior and may reveal molecular pathways relevant to renal ciliopathies.

The role of ARL13B in ciliogenesis appears to vary by cell-type. In several tissues, as well as mouse embryonic fibroblasts, loss of ARL13B or loss of ciliary ARL13B reduces ciliation and ciliary length (Caspary et al. 2007; Larkins et al. 2011; Gigante et al. 2020). ARL13B’s role in kidney cells is distinct. Loss of ARL13B in kidney epithelial cells results in loss of cilia, whereas *Arl13b^A/A^* kidneys keep their primary cilium intact across all nephron segments and maintain normal cilia length (Van Sciver et al. 2023; Seixas et al. 2016; Li et al. 2016). These observations suggest that ARL13B regulates ciliary biology through distinct mechanisms in renal epithelia compared with other cell types. Consequently, the kidney provides a valuable model for dissecting localization-dependent functions of ARL13B.

Here, we generated two independent *Arl13b^A/A^* kidney epithelial cell lines along with matched rescue cell lines. Using RNA sequencing across mutant-rescue pairs, we defined transcriptional programs associated with the loss of ciliary ARL13B while controlling for clone-specific and genome-editing-related effects. Differentially expressed genes were enriched for pathways related to ciliary biology and kidney phenotypes, supporting the biological relevance of the dataset. These findings establish a transcriptomic framework for understanding localization-dependent ARL13B function and provide insight into how subcellular compartmentalization of a ciliary signaling regulator influences gene expression.

## Methodology

### Cell culture

Murine inner medullary collecting duct (mIMCD3; ATCC CRL-2123) cells were maintained in DMEM/F12 media (Corning, 16-405-CV) supplemented with 10% fetal bovine serum (FBS) at 37°C in 5% CO . To induce ciliogenesis, cells were serum-starved in medium containing 0.5% FBS for 24 hours before analysis.

### Generation of cell lines

mIMCD3 cell lines were edited at the *Arl13b* locus using CRISPR/CAS9 to generate two independent cell lines (Synthego) using guide RNA: UAAGAAUGAAAAGGAGUCAU to cut at chr16:62,802,793 (GRCm38/mm10). Donor oligo AGGCGTGGGACTCTTTGGAGTAGACT-CGTCTGTATTGACTGGTTC<u>TGC</u>aCGATGACTCCTTTTCATTCTTAGTTTCTTGGTTTTCTTCTTACT was used to modify valine (GTA) to alanine (GCA) at <u>amino acid 358</u> of ARL13B, with a conserved base change in amino acid 357 (CGG to CGT, both coding for arginine) introduced to destroy the PAM site and prevent recutting. Individual clones were isolated by limiting dilution and proper editing was confirmed by PCR and Sanger sequencing, using primers located outside of the donor sequence: fwd-TACTGCGTAAAGCTATGCTAATACA and rev-ATGCTTTTGGTCATTTGTGTTTCA.

For each ARL13B^A/A^ clone (1-ARL13B^A^ and 2-ARL13B^A^), stable lines expressing wild-type *ARL13B-V5-His* were generated to act as rescue cell lines. Plasmid pDEST40-Arl13b11 (Addgene #40873) was linearized using XbaI and transfected into mIMCD3 cells in a 6-well plate. Stable populations were selected with 400ug/ml G418 antibiotic for four days. Individual clones were selected by limiting dilution in a 96-well plate. Ten rescue clones were chosen for each ARL13B^A/A^ clone and western blot and immunofluorescent staining for ARL13B were performed. The clones that had the highest ARL13B expression by western blot and ciliary staining most similar to wild-type mIMCD3s were chosen to act as the rescue cell lines (1R-ARL13B^A^ and 2R-ARL13B^A^).

### Immunofluorescence microscopy and ciliary quantification

Cells were cultured on glass coverslips and serum starved (0.5% serum) for 24 hours before fixation with 4% paraformaldehyde. Samples were blocked in Antibody Wash Solution (1% heat-inactivated goat serum, 0.1% TritonX-100 in TBS) and incubated overnight at 4°C with primary antibodies in Antibody Wash Solution. Coverslips were washed three times for 5 min with Antibody Wash Solution at room temperature, followed by incubation with fluorophore-conjugated secondary antibodies for 1 hour at room temperature. Coverslips were washed again three times for 5 min at room temperature and mounted using ProLong Gold Antifade Mountant (Thermo Fisher Scientific). Antibodies used: mouse anti-acetylated-alpha-tubulin (1:2500, Sigma T6793), rabbit anti-ARL13B (1:500, ProteinTech 17711-1-AP), rabbit anti-ARL3 (1:2000, (Cavenagh et al. 1994)), rabbit anti-INPP5E (1:150, ProteinTech 17797-AP), Hoechst (1:5000, Thermofisher 33342), goat anti-mouse-AF647 (1:500, Jackson Immunology 115-606-003), and donkey anti-rabbit-488 (1:500, Invitrogen A31572).

Images were acquired using a BioTek Lionheart FX automated fluorescence microscope. Z-stack images were collected at 60x with 0.5 µm intervals for cilia length measurements, whereas single focal plane images were used for ciliation frequency and fluorescence intensity analyses.

Ciliary measurements were performed using Fiji/ImageJ 2.16.0/1.54p (Schindelin et al. 2012). Fluorescent intensities were measured using the CiliaQ plugin (v0.1.7) (Burgdorf et al. 2025). Cilia were identified by staining for acetylated alpha-tubulin and a mask was obtained using a background subtraction of rolling radius 10 and using Renyi Entropy thresholding. To determine mean cilium length, measurements were made for three technical replicates of 100 cilia each (N=300). The percentage of ciliated cells was measured by counting the number of cells and ciliated cells in an image (excluding the cells touching the x,y borders) totaling three technical replicates of 100 cilia each (N=300). For fluorescence intensity analysis, the mean intensity of the brightest 10% of pixels within the ciliary mask was measured for the protein of interest. This was used as the fluorescence measurement because this metric is less sensitive to mask boundary effects and background noise than whole cilium mean intensity and more accurately reflects protein enrichment within a cilium. To further correct for background fluorescence, the mean intensity of a region of interest (ROI) adjacent to the cilium was subtracted from the ciliary measurement, yielding a background-corrected intensity value. Background-corrected fluorescence intensities were plotted as violin plots and are reported in arbitrary units (a.u.). Within each plot, the dashed lines represent the median and interquartile range. Significance between samples was measured via nested one-way ANOVA. All representative images were identically adjusted for brightness and contrast within an experiment.

### Western blot analysis

Whole-cell lysates were prepared from confluent 10cm cultures using Pierce RIPA buffer (Thermo Scientific 89901) supplemented with protease inhibitors (Thermo Scientific A32955). Protein concentrations were determined using a bicinchoninic acid (BCA) assay, and equal amounts of protein (25 ug) were separated by 10% SDS-PAGE before transfer to nitrocellulose membranes at 30v overnight at 4°C. To check for consistent protein loading and transfer, blots were Ponceau stained (VWR 102876-972). After blocking with Pierce Protein-Free Blocking Buffer (Thermo Scientific 37572), membranes were incubated with primary antibodies diluted in the blocking buffer for two hours at room temperature. The membrane was then washed three times for ten minutes each using TBST (20 mM Tris, 150 mM NaCl, 0.1% (w/v) Tween 20 detergent in water) and fluorescent secondary antibodies diluted in blocking buffer were incubated for one hour at room temperature. The membrane was washed with TBST again three times for ten minutes each. Proteins were visualized using an Azure 600 imaging system. Antibodies used: rabbit anti-ARL13B (1:1000, ProteinTech 17711-1-AP), mouse anti-HSP-90 (1:5000, Santa Cruz SC-13119), donkey anti-mouse-680RD (1:5000, LICOR 926-68072), donkey anti-rabbit-800CW (1:5000, LICOR 925-32213). Relative ARL13B expression was measured using Fiji/ImageJ 2.16.0/1.54p normalized to HSP-90 as a control (Schindelin et al. 2012).

### RNA sequencing and analysis

Three replicates of each cell line were expanded for two passages and serum-starved for 24 hours to induce ciliation before RNA isolation. Total RNA was extracted using the RNeasy Mini Kit (Qiagen) with on-column DNase digestion according to the manufacturer’s instructions. RNA concentration and purity were assessed spectrophotometrically prior to library preparation to meet Novogene standards (OD260/280 ≥ 2.0 by NanoDrop). Sequencing libraries were prepared and sequenced by Novogene using paired-end 150-bp Illumina NovaSeq 6000 sequencing. Raw sequencing reads were processed using the Galaxy platform (Galaxy 2026). Read quality was assessed using FastQC v0.12.1 (Galaxy version 0.73+galaxy0). Reads were aligned to the mouse reference genome (GRCm39/mm39) using RNA STAR v2.7.11b (Galaxy Version 2.7.11b+galaxy1), and gene-level counts were generated using featureCounts v2.0.3 (Galaxy Version 2.1.1+galaxy1) (Dobin et al. 2013; Liao et al. 2014).

Differential gene expression analysis was performed using EdgeR v3.42.4 (Galaxy version 3.42.4+galaxy0) (Robinson et al. 2010). The experimental design was treated as a single-factor layout containing three levels: WT, Mutant, and Rescue. Sample libraries were normalized using the Trimmed Mean of M-values (TMM) method. Biological variation and dispersion parameters were estimated using the empirical Bayes method. Pairwise group comparisons were evaluated through three specific contrasts (Mutant vs. WT, Rescue vs. WT, and Mutant vs. Rescue) using edgeR’s quasi-liklihood (QL) F-test framework. P-values were adjusted for multiple testing using the Benjamini-Hochberg false discovery rate (FDR) procedure, with significance defined at an FDR threshold of < 0.05 and genes with a |logFC| > 1 were considered significantly differentially expressed. Principal component analysis plots were generated using ggplot2 v3.5.1 (Galaxy version 3.5.1+galaxy2) and heatmaps were generated using heatmap2 (Galaxy Version 3.3.0+galaxy0) (Wickham 2009; heatmap.2: Enhanced Heat Map). Gene enrichment analysis was performed using the GOEnrichment tool in Galaxy (Galaxy version 2.0.1+galaxy0) using all expressed genes as the background set (Faria 2017). Significantly enriched terms (P < 0.05) were used for downstream biological interpretation of differentially expressed gene sets. Renal/urinary system-related phenotypes were identified using the Jax Human Mouse Disease Connection (RRID:SCR_017522) Portal (Eppig et al. 2015). Overlap of DEGs with regulatory target gene sets was queried using the M3 collection (regulatory target gene sets) in the Molecular Signatures Database (MSigDB) (Castanza et al. 2023; Subramanian et al. 2005).

### Statistical analysis

Statistical analyses were performed using GraphPad Prism 11.0.0. Unless otherwise stated, data are presented as mean ± SEM. Comparisons among multiple groups were performed using nested one-way ANOVA followed by appropriate post hoc multiple-comparison tests. Statistical significance was defined as P < 0.05.

## Results

### Loss of ciliary ARL13B in mIMCD3 cells does not alter ciliogenesis

To investigate the function of ciliary ARL13B in kidney cells, we established two mIMCD3 cell lines expressing only ARL13B^A^ from the endogenous locus using CRISPR/CAS9 (see methods for details). We refer to control mIMCD3 cells that went through CRISPR without the guide as WT and to the individual clones as 1-ARL13B^A^ and 2-ARL13B^A^ (Figure 1A). To control for off-target effects, we generated stable *Arl13b-V5-His* expressing rescue lines for each clone. Using western blot and immunofluorescent (IF) staining, we identified a robust expressing rescue line for each clone for subsequent analysis; we call these clones 1R- ARL13B^A^ and 2R-ARL13B^A^ respectively. Through western blot analysis, we found ARL13B expression across all cell lines (Figure 1B, Supplemental Figure 1).

**Figure 1:**
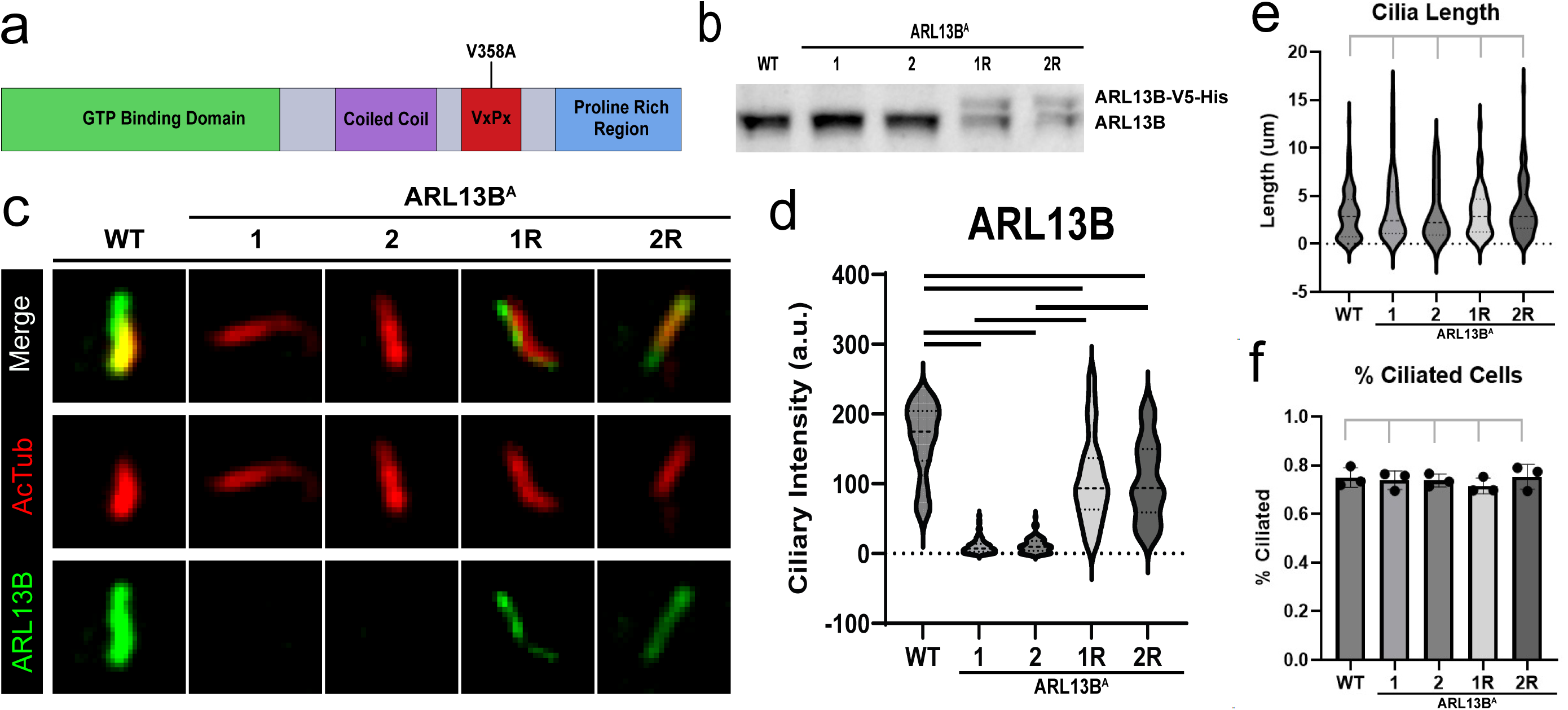
ARL13B^A^ is undetectable in cilia and does not impact ciliogenesis in mIMCD3 cells. (A) Schematic of ARL13B domains indicating the V358A mutation location in the ciliary localization signal. (B) Western blot on all cell lines detecting ARL13B. (C) Immunofluorescent images of mIMCD3 cell lines showing localization of ARL13B (green) to cilia (AcTub, red). (D) Quantification of corrected ciliary fluorescent intensity of ARL13B. Three technical replicates of 30 cilia each, total N=90. (E) Quantification of cilia length (um) of mIMCD3 cell lines. Three technical replicates of 100 cilia each, total N=300. (F) Percent of ciliated cells (cilia/nucleus) in mIMCD3 cell lines. Three technical replicates of 100 cilia each, total N=300. All analyses underwent nested one-way ANOVA significance testing (p<0.05). Black bars across comparisons indicate significant differences (p<0.05) and gray bars across comparisons indicate no significant difference. Alt text: Figure 1 demonstrates that the ARL13B^V358A^ mutation excludes ARL13B from primary cilia without affecting ciliogenesis. A schematic shows the V358A mutation within the ciliary localization sequence of ARL13B, immunofluorescence images show ciliary ARL13B present in wild-type and rescue cells but absent from mutant cells, and quantification confirms a significant reduction in ciliary ARL13B signal in both mutant clones. Western blot analysis shows ARL13B expression across cell lines. Measurements of cilia length and the percentage of ciliated cells reveal no significant differences among wild-type, mutant, and rescue lines, indicating that loss of ciliary ARL13B does not impair ciliogenesis in mIMCD3 cells.

We examined ARL13B localization using IF staining of ARL13B and the ciliary marker, acetylated α-tubulin (AcTub). We observed ARL13B enrichment in cilia of WT cells as well as in rescue cell lines 1R-ARL13B^A^ and 2R-ARL13B^A^. In contrast, we detected no ciliary ARL13B in either 1-ARL13B^A^ or 2-ARL13B^A^ lines, consistent with the ARL13B^A^ mutation disrupting the ciliary enrichment of ARL13B (Gigante et al. 2020; Mariani et al. 2016) (Figure 1C). We quantified the staining intensity of ciliary ARL13B and found both WT and 1R&2R-ARL13B^A^ cell lines displayed significantly more ciliary ARL13B than 1&2-ARL13B^A^ cell lines. 1R&2R-ARL13B^A^ cell lines exhibited less ciliary ARL13B than WT, consistent with the western analysis (Figure 1D). These data argue that the ARL13B^A^ CRISPR-induced mutation disrupts the ciliary localization of ARL13B and that we can rescue ciliary localization through ARL13B-V5-His stable expression.

As ARL13B regulates ciliogenesis in kidney cells in a distinct manner compared to other cell types, we quantified the percentage of ciliated cells as well as the length of the cilia in the WT, 1-ARL13B^A^ ,1R-ARL13B^A^, 2-ARL13B^A^ and 2R-ARL13B^A^ mIMCD3 cell lines. We found no significant difference in average cilia length between cell types, with an average cilia length of 3.26 + 0.24 um (p>0.05) (Figure 1E). Additionally, we found the average percentage of cells ciliated after 24 hours of serum starvation was 73.86% + 0.015% with no significant difference among genotypes (p>0.05) (Figure 1F). These results indicate that loss of ciliary ARL13B does not impact ciliogenesis in mIMCD3 cell lines.

### Cilia protein composition is impacted by loss of ciliary ARL13B

We examined the localization of several proteins known to be mislocalized in the absence of ARL13B. We detected ciliary INPP5E in WT mIMCD3s but not in 1&2-ARL13B^A^ lines (Figure 2A). Similarly, we found ciliary ARL3 staining in WT mIMCD3s which was lost in 1&2-ARL13B^A^ cells (Figure 2B). Of note, both INPP5E and ARL3 staining in 1R&2R-ARL13B^A^ cell lines was significantly more intense than in 1&2-ARL13B^A^ cells and significantly less intense than in WT, suggesting a partial rescue (Figure 2B).

**Figure 2:**
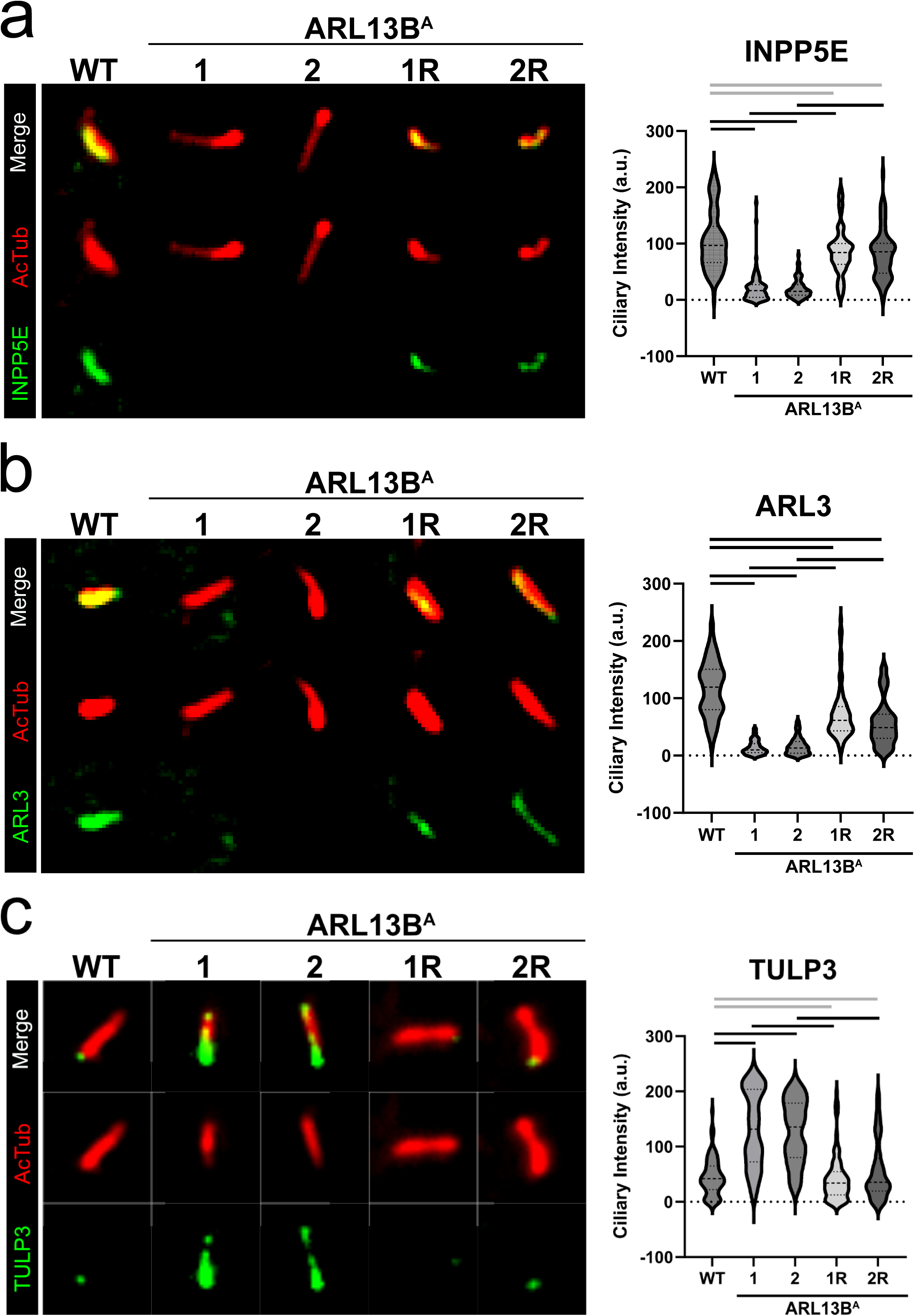
Changes in localization of ciliary proteins as a result of ARL13B^A^. Quantification of corrected ciliary fluorescent intensity of INPP5E (A), ARL3 (B), TULP3 (C). Three technical replicates of 30 cilia each, total N=90. Representative images display localization of the indicated protein of interest (green) to cilia (AcTub, red). All analyses underwent nested one-way ANOVA significance testing (p<0.05). Black bars across comparisons indicate significant differences (p<0.05) and gray bars across comparisons indicate no significant difference. Alt text: Immunofluorescence images and quantitative analysis of ciliary protein localization in WT, 1&2-ARL13B^A^, and 1R&2R-ARL13B^A^ cell lines. Cilia are marked by acetylated tubulin (red) and the indicated protein (green). INPP5E and ARL3 show strong ciliary localization in WT cells, reduced localization in ARL13B^A^ cells, and restoration in rescue lines. TULP3 shows increased ciliary localization in ARL13B^A^ cells that is reduced in rescue lines. Quantification of 90 cilia per condition confirms significant changes in INPP5E, ARL3, and TULP3 localization following ARL13B loss, with rescue restoring localization toward WT levels.

In contrast, we observed significantly more ciliary TULP3 staining in 1&2-ARL13B^A^ cell lines than in WT controls (Figure 2C). TULP3 ciliary staining intensity in 1R&2R-ARL13B^A^ cell lines was indistinguishable from WT (Figure 2C). These results indicate that ARL13B is required in cilia for the proper localization of ARL3, INPP5E, and TULP3 to the cilium in mIMCD3 cell lines.

### Transcriptional effect of loss of ciliary ARL13B

To determine the impact on transcription from the loss of ciliary ARL13B in mIMCD3 cells, we performed bulk RNAseq on WT, 1&2-ARL13B^A^ and 1R&2R-ARL13B^A^ cell lines (Figure S1). We performed two distinct, pairwise differential expression analyses to assess the specific impact of ciliary ARL13B. We grouped 1-ARL13B^A^ and 2-ARL13B^A^ to compare to WT (1&2-ARL13B^A^ vs WT) and identified 631 differentially expressed genes (DEGs, adjusted *P* < 0.05, |logFC| > 1) (Fig 3A; Table S1). We also combined rescue clones 1 and 2 to compare to WT (1R&2R-ARL13B^A^ vs WT) and found 380 DEGs (Fig 3B). Of the 631 DEGs identified in the 1&2-ARL13B^A^ vs WT comparison, 417 were rescued in the 1R&2R-ARL13B^A^ cell lines, no longer meeting the threshold for differential expression (Figure 3B). We focused on these 417 genes because their expression changes are attributable to the loss of ciliary ARL13B function; hereafter, they are referred to as the 1&2-ARL13B^A^ vs WT cilia-specific DEGs (Figure 3C). The 417 DEGs exhibited varying levels of expression in 1R&2R-ARL13B^A^ compared to 1&2-ARL13B^A^ (Figure 3D). Through gene ontology enrichment analysis, we found DEGs related to processes including cilia function and structure (establishment of left/right asymmetry, axonemal B tubule inner sheath), mechanotransduction (cellular response to fluid shear stress, integrin alphav-beta3 complex), and epithelial morphogenesis (morphogenesis of an epithelium, desmosome, protein complex involved in cell adhesion) (Figure 3E; Table S2). Together, these data provide a transcriptomic signature specific to loss of ciliary ARL13B in kidney cells.

**Figure 3:**
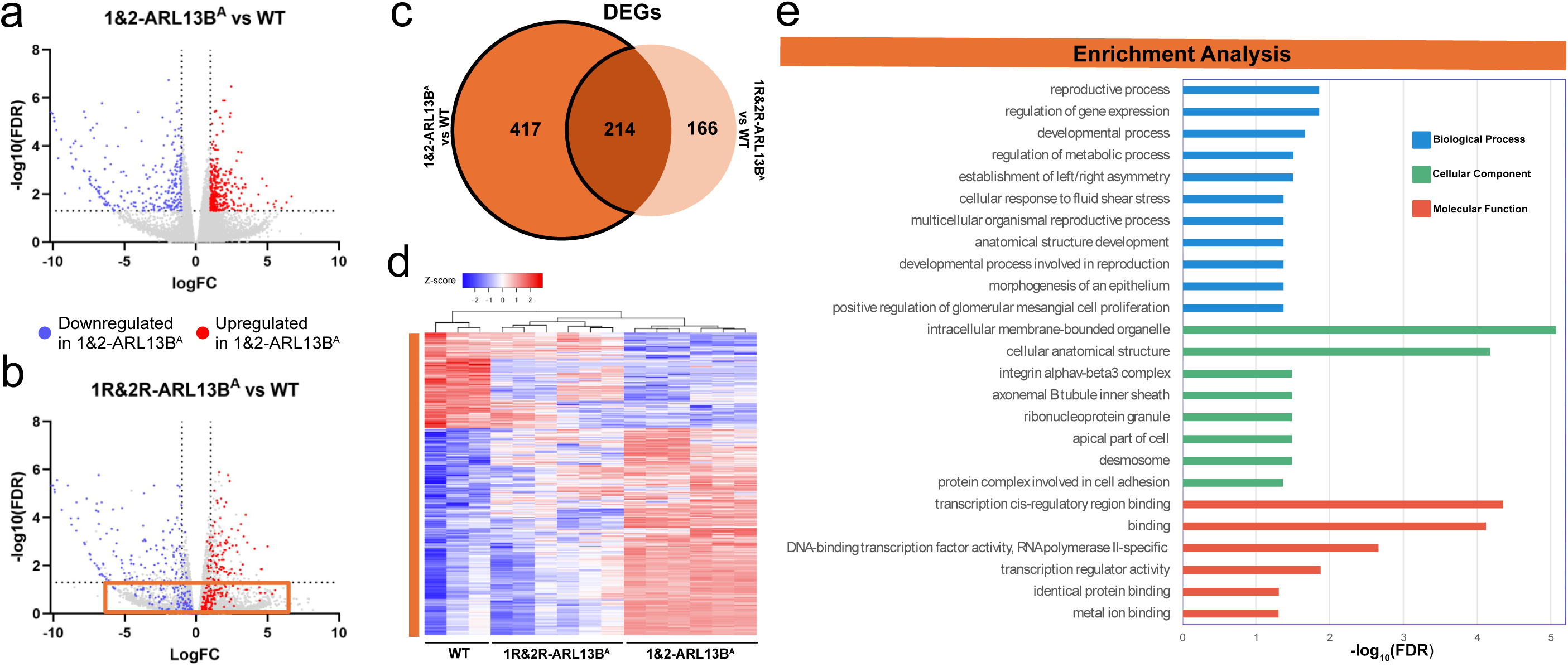
1&2-ARL13B^A^ transcriptomics. (A) Volcano plot of 1&2-ARL13B^A^ vs WT pairwise comparison. (B) Volcano plot of 1R&2R-ARL13B^A^ vs WT pairwise comparison. Differentially expressed genes (|logFC| > 1, adjusted *P* < 0.05) found in the 1&2-ARL13B^A^ vs WT comparison are shown in blue (downregulated) and red (upregulated) for both A and B. DEGs from the first comparison rescued in 1R&2R-ARL13B^A^ are boxed in orange. (C) Venn diagram displaying the number of DEGs in each comparison including genes overlapping between samples. The outlined portion highlights the 1&2-ARL13B^A^ vs WT ciliary ARL13B specific DEGs shown in the heatmap, gene ontology enrichment analysis, and Fig 6 analyses. (D) Heatmap of cilia-specific DEGs with Z-scores ranging from -2 (blue) to 2 (red). (E) Differentially expressed genes enrichment analysis for biological processes (blue), cellular component (green), and molecular function (red). Alt text: Figure 3 summarizes transcriptomic differences between ARL13B^A^ mutant and rescue relative to wild-type cells. Volcano plots show significant upregulated and downregulated genes in mutant cells, a Venn diagram identifies 214 shared and 417 mutant-specific differentially expressed genes, a heatmap demonstrates distinct expression patterns in mutant versus WT and rescue cells, and gene ontology analysis reveals enrichment of developmental, cilia-related, cell adhesion, and transcriptional regulatory pathways.

### Clonal differences result in distinct transcriptomic profiles

To ascertain the similarities and distinctions between the 1-ARL13B^A^ and 2-ARL13B^A^ cell lines, we repeated the two pairwise differential expression analyses on the 1-ARL13B^A^ and 2-ARL13B^A^ cell lines individually. For 1-ARL13B^A^, we identified 1870 DEGs (1-ARL13B^A^ vs. WT; adjusted *P* < 0.05, |logFC| > 1) (Figure 4A; Table S3). We found 2229 DEGs in the 1R-ARL13B^A^ vs WT comparison. Among the 1870 1-ARL13B^A^ vs WT DEGs, we identified 1249 (1-ARL13B^A^ vs WT cilia-specific DEGs) that were rescued in 1R-ARL13B^A^ (falling below the significance threshold), albeit to varying levels (Figure 4B-D). Using gene ontology enrichment analysis, we observed DEGs relating to pathways involved in cilia function (ciliary basal body, establishment of left/right asymmetry, structural constituent of cytoskeleton) and tissue remodeling (response to transforming growth factor beta, negative regulation of angiogenesis), among others (Figure 4E; Table S2).

**Figure 4:**
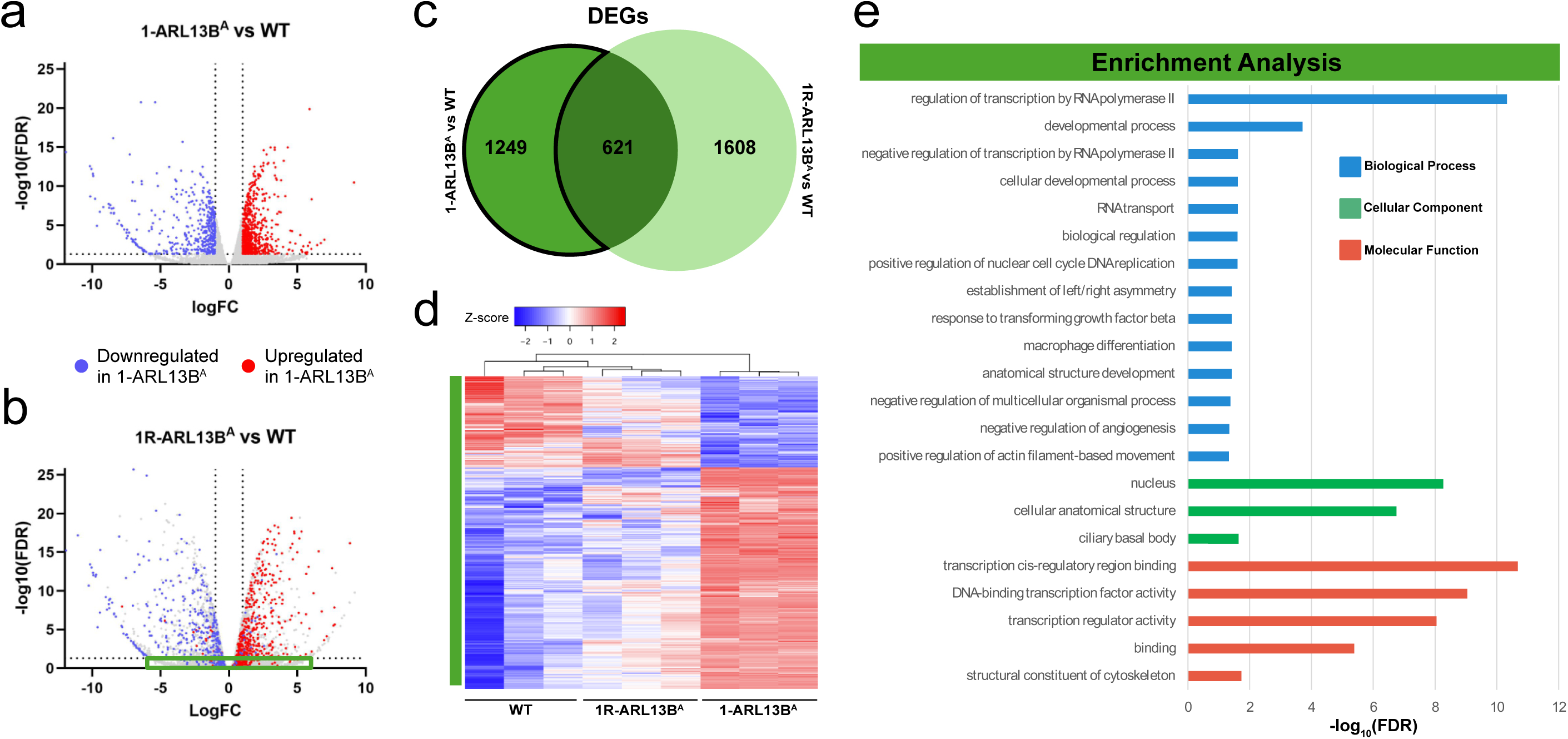
1-ARL13B^A^ transcriptomics. (A) Volcano plot of 1-ARL13B^A^ vs WT pairwise comparison. (B) Volcano plot of 1R-ARL13B^A^ vs WT pairwise comparison. Differentially expressed genes (|logFC| > 1, adjusted *P* < 0.05) found in the 1-ARL13B^A^ vs WT comparison are shown in blue (downregulated) and red (upregulated) for both A and B. DEGs from the first comparison rescued in 1R-ARL13B^A^ are boxed in green. (C) Venn diagram displaying the number of DEGs in each comparison, including genes overlapping between samples. The outlined portion reflects the 1-ARL13B^A^ vs WT ciliary ARL13B specific DEGs shown in the heatmap, gene ontology enrichment analysis, and Fig 6 analyses. (D) Heatmap of cilia-specific DEGs with Z-scores ranging from -2 (blue) to 2 (red). (E) Differentially expressed genes enrichment analysis for biological processes (blue), cellular component (green), and molecular function (red). Alt text: Figure 4 summarizes transcriptomic changes in 1-ARL13B^A^ mutant and rescue cells relative to wild type. Volcano plots identify extensive upregulated and downregulated genes in mutant cells, a Venn diagram shows 621 shared DEGs and 1,249 mutant-specific DEGs, a heatmap demonstrates distinct expression patterns in mutant cells compared with WT and rescue samples, and gene ontology analysis reveals enrichment of transcriptional regulation, developmental pathways, nuclear and ciliary components, and cytoskeletal functions.

For the 2-ARL13B^A^ clone analysis, we identified 1618 DEGs (2-ARL13B^A^ vs WT; adjusted *P* < 0.05, |logFC| > 1) (Figure 5A; Table S4). We found 1336 differentially expressed genes for 2R-ARL13B^A^ (2R-ARL13B^A^ vs WT), with 1120 of the 1618 genes differentially expressed in the 2-ARL13B^A^ vs WT comparison falling below the significance threshold in the rescue (Figure 5B,C). There were varying levels of rescue in the 1120 cilia-specific DEGs (2-ARL13B^A^ vs WT) (Figure 5D). We observed DEGs relating to pathways involved in tubular metabolism (oxidoreductase activity, acting on NAD(P)H), epithelial structure (integrin complex, apical plasma membrane, anchoring junction, plasma membrane protein complex), and signal transduction (signaling receptor complex, cell surface receptor signaling pathway, regulation of signal transduction) (Figure 5E; Table S2). Together, these individual cell line analyses (1-ARL13B^A^ and 2-ARL13B^A^) indicate that many differences exist between the clones both in the category of rescued DEGs (likely on-target) and non-rescued (potential clonal effect).

**Figure 5:**
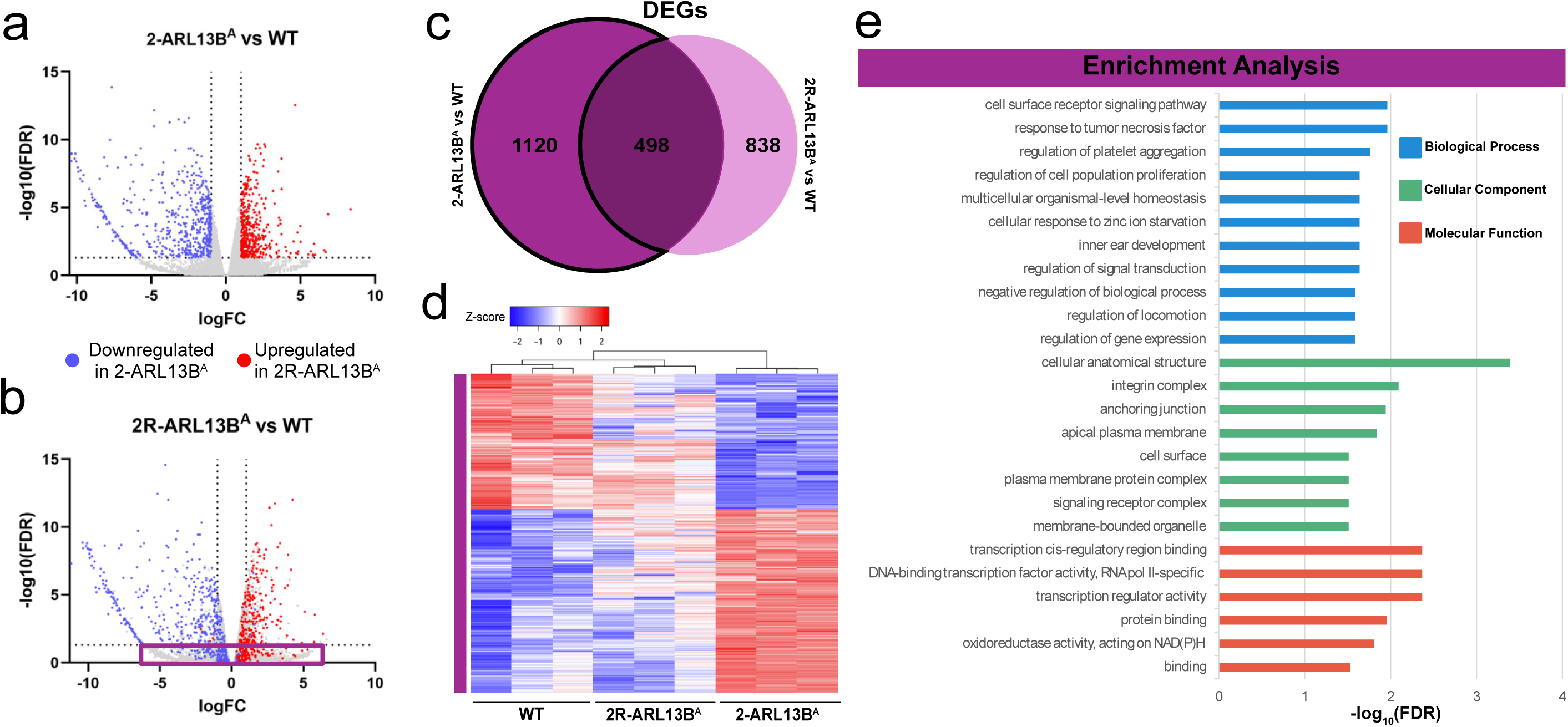
2-ARL13B^A^ transcriptomics. (A) Volcano plot of 2-ARL13B^A^ vs WT pairwise comparison. (B) Volcano plot of 2R-ARL13B^A^ vs WT pairwise comparison. Differentially expressed genes (|logFC| > 1, adjusted *P* < 0.05) found in the 2-ARL13B^A^ vs WT comparison are shown in blue (downregulated) and red (upregulated) for both A and B. DEGs from the first comparison rescued in 2R-ARL13B^A^ are boxed in purple. (C) Venn diagram displaying the number of DEGs in each comparison including genes overlapping between samples. The outlined portion signifies the 2-ARL13B^A^ vs WT ciliary ARL13B specific DEGs shown in the heatmap, gene ontology enrichment analysis, and Fig 6 analyses. (D) Heatmap of cilia-specific DEGs with Z-scores ranging from -2 (blue) to 2 (red). (E) Differentially expressed genes enrichment analysis for biological processes (blue), cellular component (green), and molecular function (red). Alt text: Figure 5 summarizes transcriptomic changes in 2-ARL13B^A^ mutant and rescue cells relative to wild type. Volcano plots identify significantly upregulated and downregulated genes in mutant cells, a Venn diagram shows 498 shared DEGs with 1,120 mutant-specific and 838 rescue-specific DEGs, a heatmap demonstrates genotype-dependent expression patterns that distinguish mutant from WT and rescue samples, and gene ontology analysis reveals enrichment of cell signaling, gene regulation, cell surface and membrane-associated components, integrin complexes, and transcription-related molecular functions.

### Prioritization of ciliary ARL13B DEGs

While all DEGs identified in the 1&2-ARL13B^A^, 1-ARL13B^A^, or 2-ARL13B^A^ vs WT comparisons are potential targets, any DEGs we found that were rescued in the 1R&2R-ARL13B^A^, 1R-ARL13B^A^ or 2R-ARL13B^A^ vs WT analyses are targets specific to loss of ciliary ARL13B and can be prioritized for follow up studies (hereafter called cilia-specific DEGs) (Fig 6). We reasoned that we could further prioritize a subset of DEGs by identifying those in common among all three analyses. Thus, we evaluated the 417 cilia-specific DEGs from the 1&2-ARL13B^A^ vs WT comparison, the 1249 cilia-specific DEGs from the 1-ARL13B^A^ vs WT comparison, and the 1120 cilia-specific DEGs from the 2-ARL13B^A^ vs WT comparison; we found 131 overlapping DEGs (Figure 6A, Table S5). Among the 131 genes, we identified ontology terms including those related to ciliogenesis, left/right asymmetry, and macrophage differentiation (Figure 6B). Additionally, several of the differentially expressed genes were found to have mammalian phenotypes related to kidney function and disease (Table 1). This includes *Ccdc39*, a gene encoding a ciliary axonemal protein whose mutation is a known cause of primary ciliary dyskinesia (PCD), and *Jag1*, which is notable due to its established functions in renal epithelial differentiation and Notch signaling during kidney development (Cheng et al. 2007; Merveille et al. 2011; McCright et al. 2002). To identify potential upstream transcriptional and post-transcriptional regulators driving the observed expression changes, we performed an overrepresentation analysis on the 131 ciliary ARL13B specific DEG dataset. Notably, the dataset showed a significant enrichment for OGT target genes (FDR = 4.98E-02) and several microRNAs (Table 2).

**Figure 6:**
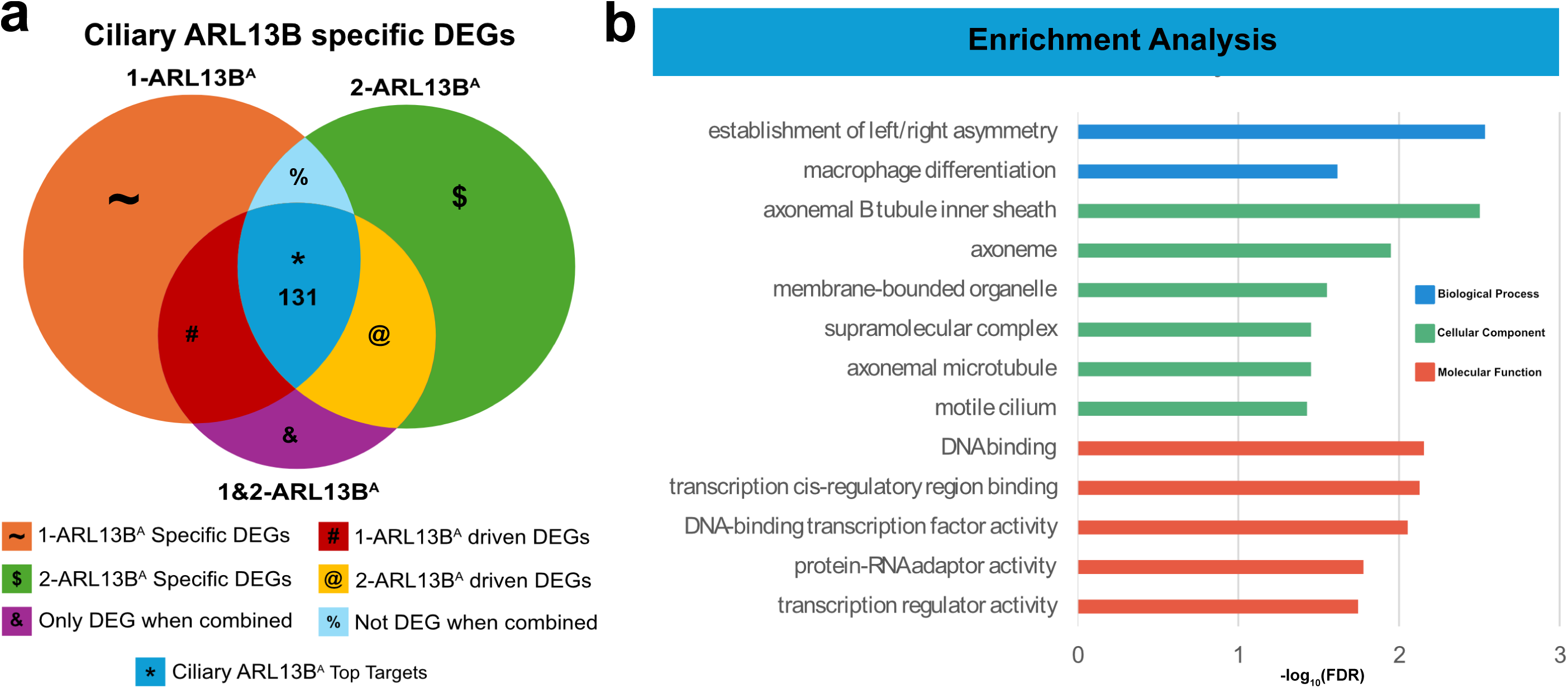
Prioritization of ciliary ARL13B-dependent transcriptional targets. (A) Venn diagram comparing ciliary ARL13B-specific differentially expressed genes (DEGs) identified in the 1-ARL13B^A^, 2-ARL13B^A^, and combined 1&2-ARL13B^A^ analyses. The overlap of all three analyses identified 131 high-confidence ciliary ARL13B-dependent targets (*, blue). Additional regions indicate clone-specific, clone-driven, and combined-analysis-dependent DEGs. (B) Gene ontology enrichment analysis of the 131 overlapping high-confidence targets identified in panel A. Enriched biological process terms are shown in blue, cellular component terms in green, and molecular function terms in red. Alt text: Figure 6 presents a Venn diagram comparing ciliary ARL13B-dependent differentially expressed genes identified in the 1-ARL13B^A^, 2-ARL13B^A^, and combined 1&2-ARL13B^A^ analyses. The overlap among all three analyses contains 131 high-confidence targets, while additional regions represent clone-specific, clone-driven, and combined-analysis-dependent gene sets. A gene ontology enrichment analysis of the 131 shared targets shows enrichment for cilia-related and developmental processes, including left-right asymmetry and axonemal structures, as well as transcriptional regulatory functions.

**Table 1:**
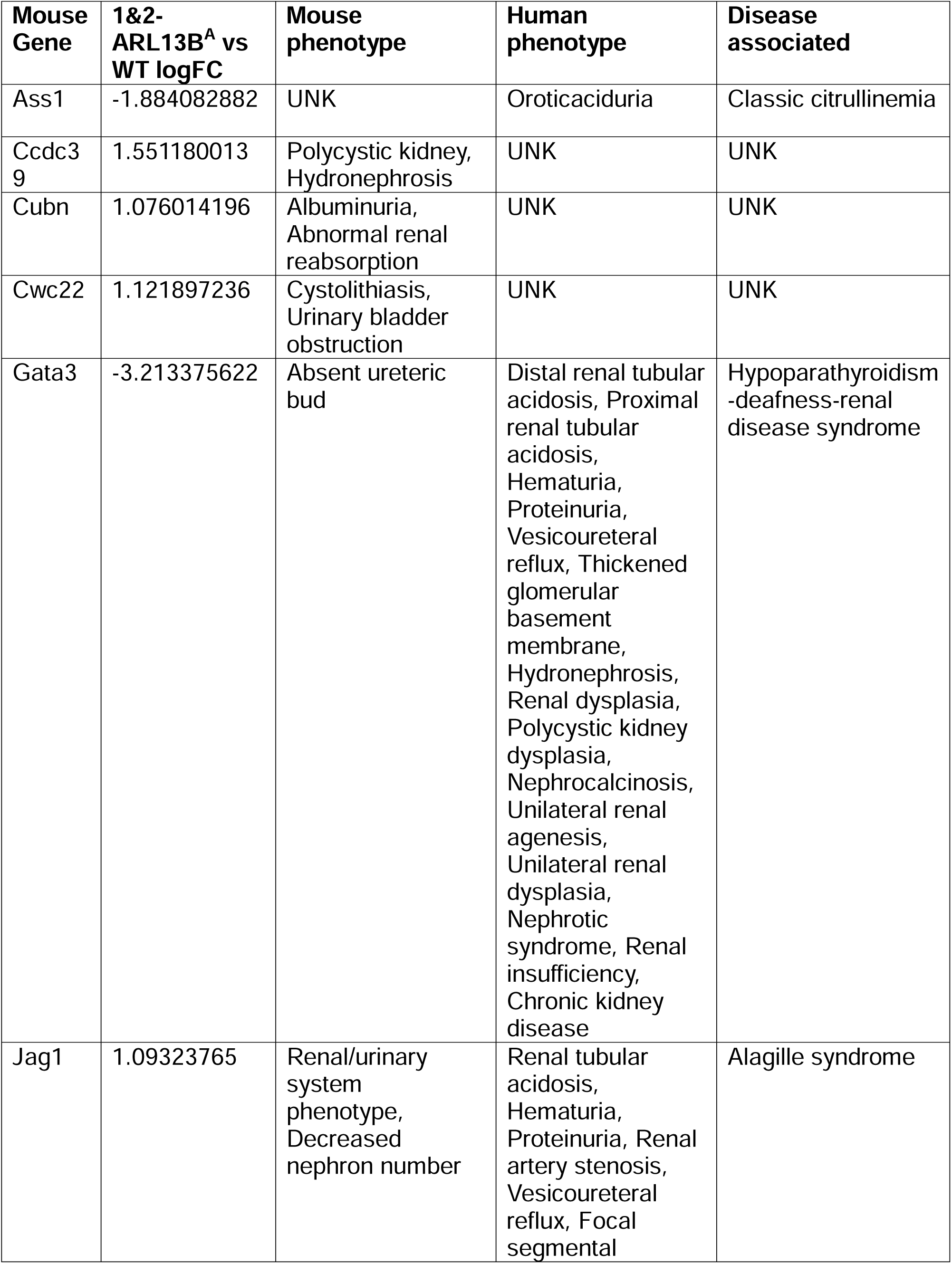
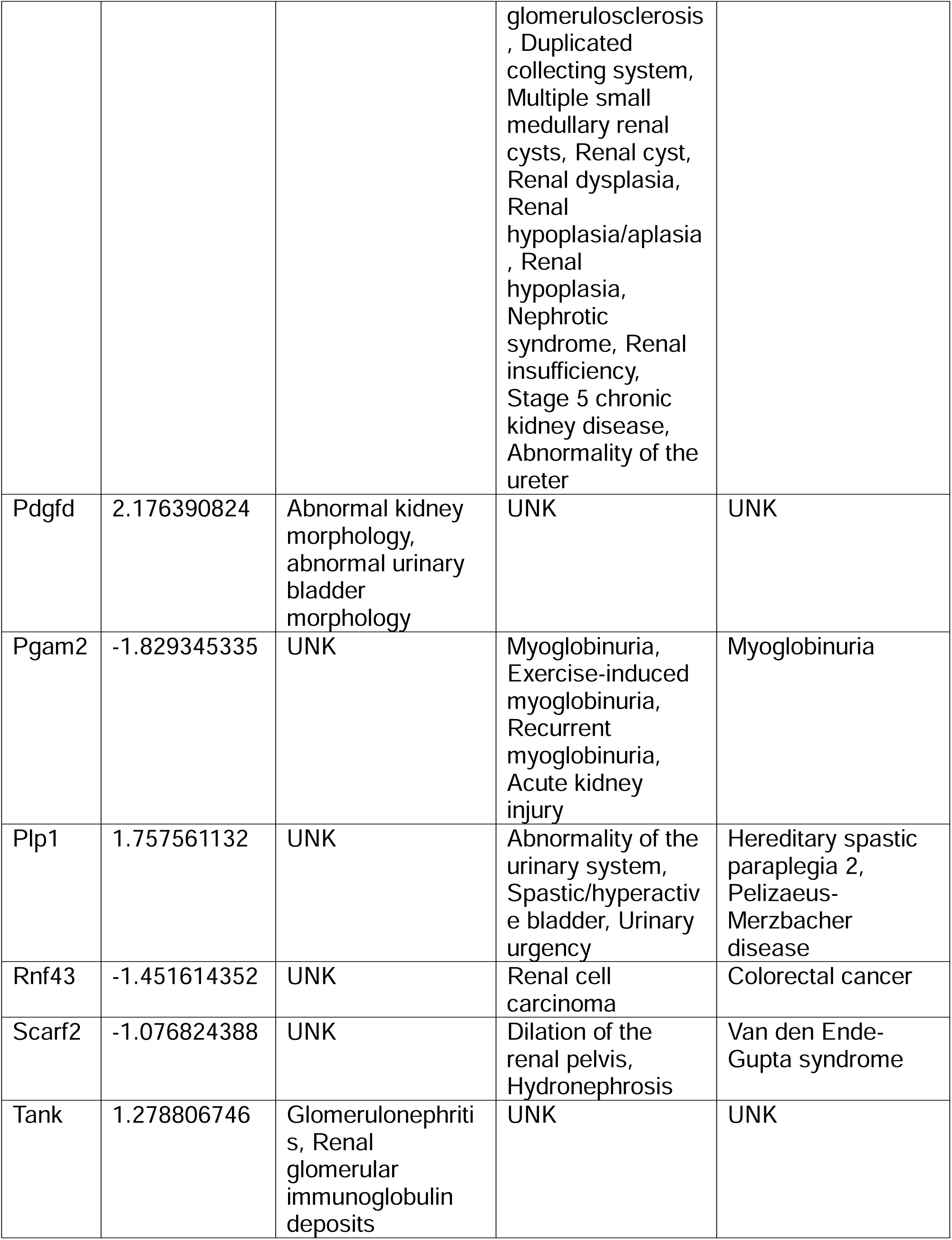
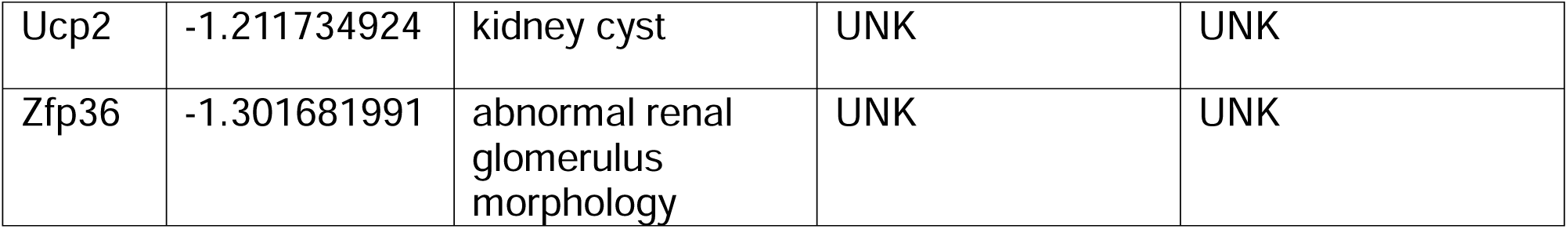
Targets with kidney phenotypes. Table of genes found in the ciliary-specific ARL13B DEGs that have known mouse kidney phenotypes (column 3), human kidney phenotypes (column 4) and or human disease associated (column 5). LogFC of DEGs is listed in column 2 and unknown phenotypes depicted as UNK.

| **Mouse Gene** | **1&2-ARL13B<sup>A</sup> vs WT logFC** | **Mouse phenotype** | **Human phenotype** | **Disease associated** |
| --- | --- | --- | --- | --- |
| Ass1 | -1.884082882 | UNK | Oroticaciduria | Classic citrullinemia |
| Ccdc39 | 1.551180013 | Polycystic kidney, Hydronephrosis | UNK | UNK |
| Cubn | 1.076014196 | Albuminuria, Abnormal renal reabsorption | UNK | UNK |
| Cwc22 | 1.121897236 | Cystolithiasis, Urinary bladder obstruction | UNK | UNK |
| Gata3 | -3.213375622 | Absent ureteric bud | Distal renal tubular acidosis, Proximal renal tubular acidosis, Hematuria, Proteinuria, Vesicoureteral reflux, Thickened glomerular basement membrane, Hydronephrosis, Renal dysplasia, Polycystic kidney dysplasia, Nephrocalcinosis, Unilateral renal agenesis, Unilateral renal dysplasia, Nephrotic syndrome, Renal insufficiency, Chronic kidney disease | Hypoparathyroidism -deafness-renal disease syndrome |
| Jag1 | 1.09323765 | Renal/urinary system phenotype, Decreased nephron number | Renal tubular acidosis, Hematuria, Proteinuria, Renal artery stenosis, Vesicoureteral reflux, Focal segmental glomerulosclerosis , Duplicated collecting system, Multiple small medullary renal cysts, Renal cyst, Renal dysplasia, Renal hypoplasia/aplasia , Renal hypoplasia, Nephrotic syndrome, Renal insufficiency, Stage 5 chronic kidney disease, Abnormality of the ureter | Alagille syndrome |
| Pdgfd | 2.176390824 | Abnormal kidney morphology, abnormal urinary bladder morphology | UNK | UNK |
| Pgam2 | -1.829345335 | UNK | Myoglobinuria, Exercise-induced myoglobinuria, Recurrent myoglobinuria, Acute kidney injury | Myoglobinuria |
| Plp1 | 1.757561132 | UNK | Abnormality of the urinary system, Spastic/hyperactive bladder, Urinary urgency | Hereditary spastic paraplegia 2, Pelizaeus-Merzbacher disease |
| Rnf43 | -1.451614352 | UNK | Renal cell carcinoma | Colorectal cancer |
| Scarf2 | -1.076824388 | UNK | Dilation of the renal pelvis, Hydronephrosis | Van den Ende-Gupta syndrome |
| Tank | 1.278806746 | Glomerulonephritis, Renal glomerular immunoglobulin deposits | UNK | UNK |
| Ucp2 | -1.211734924 | kidney cyst | UNK | UNK |
| Zfp36 | -1.301681991 | abnormal renal glomerulus morphology | UNK | UNK |

**Table 2:** Regulatory target gene sets overlapping with ciliary-ARL13B-specific DEGs. Table of overlapping gene sets based on gene target predictions for microRNA seed sequences and predicted transcription factor binding sites (MSigDB M3 Regulatory Gene set) (column 1). Ciliary-ARL13B specific DEGs overlapping with gene sets (column 2). False discovery rate depicting significance of enrichment (FDR < 0.05) (column 3). * Genes predicted to be targets of miRBase v22 microRNA in miRDB v6.0 with MirTarget v4 prediction scores > 80 (high confidence targets). ** Genes containing one or more binding sites for UniProt:Q8CGY8 (Ogt) in their promoter regions (TSS - 1000,+100 bp) as identified by GTRD version 20.06 ChIP-seq harmonization.

| Gene Set Name | Genes in Overlap | FDR |
| --- | --- | --- |
| MIR_1264_3P* | *Qki, Fam76b, Maml2, Golm2, Oacyl, Dclre1c, Ccdc39* | 3.54E-03 |
| MIR_335_3P* | *Qki, Fam76b, Maml2, Golm2, Oacyl, Dclre1c, Ccdc39, Zfp449, Plp1, Sema3e, Plod2, Eml5, Hnrnpa3, Zfp867* | 1.11E-02 |
| MIR_6964_3P* | *Zfp449, Plp1, Sema3e, Zfp945, Zfp558* | 1.26E-02 |
| MIR_6947_5P* | *Zfp951, Zfp976, Myef2, Cfap52* | 1.44E-02 |
| MIR_466I_3P* | *Qki, Fam76b, Maml2, Cpne8, Slc25a27, Cep70* | 2.92E-02 |
| OGT_TARGET_GENES** | *Golm2, Plod2, Napb, Jag1, Rexo5, Ccdc171, Zfp2, Pyroxd1, Lrrfip1, Gm13421, Spo11, F7300043M19Rik, Snrnp48* | 4.98E-02 |
| MIR_511_3P* | *Golm2, Zfp449, Napb, Tank* | 4.98E-02 |
| MIR_6961_3P* | *Cpne8, Jag1, Rexo5, Enkur* | 4.98E-02 |
| MIR_466L_5P* | *Golm2, Oacyl, Zfp449, Plp1, Cpne8, Napb, Aldh1l2, Peg3, Zfp606* | 4.98E-02 |

The identification of 131 overlapping DEGs across all three analyses represent high-confidence candidates whose expression depends on ciliary ARL13B. Although more stringent criteria could be applied, such as requiring full restoration of differentially expressed genes to wildtype ARL13B expression levels in the rescue lines, this would likely exclude *bona fide* ciliary ARL13B-responsive genes because rescue of ciliary ARL13B is imperfect in the rescue cell lines we analyzed. Further filtering to the 131 DEGs (mutant vs rescue) reduces the DEG list to 44, but also likely filters out ARL13B-sensitive genes (Supplementary Figure 3). We also found secondary targets for prioritization by comparing the three analyses (Table S2). The secondary targets for prioritization include the DEGs shared in two of the comparisons or in the combined analysis (1&2-ARL13B^A^) (Figure 6A). The Venn diagram depicts the DEGs driven by 1-ARL13B^A^ or 2-ARL13B^A^ as well as those that require the 1&2-ARL13B^A^ combined analysis to be significant (Figure 6A). The secondary target list also includes the DEGs that were only partially rescued in the rescue cell lines (DEGs significant in the 1&2-ARL13B^A^ vs WT, 1R&2R-ARL13B^A^ vs WT, and 1&2-ARL13B^A^ vs 1R&2R-ARL13B^A^ analyses) and genes where the expression in rescue cells surpassed that of WT (1R&2R-ARL13B^A^ vs WT analysis) (Table S6). These data suggest that distinct analyses of overlapping DEGs provide an additional level from which to prioritize DEGs for follow up studies.

## Discussion

Our findings are consistent with previous studies demonstrating context-dependent functions of ARL13B in different cell types. ARL13B deficiency impairs ciliogenesis and reduces cilia length in mouse embryonic fibroblasts and other cell types, suggesting an important role in ciliary assembly and maintenance (Caspary et al. 2007; Larkins et al. 2011; Cevik et al. 2010). In contrast, the ARL13B^A^ variant had no detectable effect on cilia formation in mIMCD3 cells. Along with evidence showing loss of ciliary ARL13B reducing cilia length in MEFs and ARL13B^A^ mice having normal kidney cilia, this indicates that kidney epithelial cells may be less dependent on ciliary ARL13B for ciliogenesis than other cell types (Van Sciver et al. 2023; Gigante et al. 2020). Despite the preservation of structurally normal cilia, loss of ciliary ARL13B disrupted localization of its known downstream effectors ARL3 and INPP5E and resulted in accumulation of TULP3 within cilia . These observations reinforce the concept that ARL13B serves as a key regulator of ciliary composition and trafficking and suggest that disruption of these processes can occur independently of overt defects in cilia assembly (Gigante and Caspary 2020).

The transcriptional changes associated with loss of ciliary ARL13B provide a potential mechanistic link between altered ciliary signaling and renal cystogenesis. Notably, mice expressing ARL13B^A^ develop cystic kidneys despite retaining renal cilia, suggesting disease pathogenesis can arise from defective ciliary signaling rather than absence of the organelle itself (Van Sciver et al. 2023). Consistent with this model, we identified differential expression of genes and enrichment of biological processes related to ciliary function, mechanotransduction, epithelial organization, and kidney-associated phenotypes. These pathways represent plausible downstream consequences of altered ciliary composition and provide candidate mechanisms through which disrupted ciliary signaling could influence epithelial behavior and promote cyst formation. Future studies will be required to determine which of these transcriptional changes are direct consequences of ARL13B-dependent signaling and which represent secondary adaptations. In particular, genes implicated in ciliopathies, such as *Ccdc39,* and genes with established roles in renal physiology and disease, including *Ucp2, Cubn*, and *Jag1,* represent promising candidates for functional investigation. Regulatory gene set enrichment analysis further suggested that a subset of the observed transcriptional changes may be mediated through specific regulatory programs. The upregulated DEGs showed significant enrichment for promoter binding sites of OGT, as well as for targets of several microRNAs. While the pathways connecting ciliary ARL13B to these regulatory networks remain unclear, these findings identify potential downstream regulatory targets and provide new avenues for future investigation.

An important observation that emerged from this work was the substantial transcriptomic heterogeneity between independently derived CRISPR-edited clones. Although both ARL13B^A^ clones displayed highly similar ciliary phenotypes, each contained a large set of DEGs that was not shared with the other. These findings highlight a well-recognized challenge in CRISPR-based functional studies, where clonal variation, compensatory cellular responses, and unintended editing-associated effects can confound interpretation of transcriptomic data (Kosicki et al. 2018). Had either clone been analyzed in isolation, numerous genes would likely have been misclassified as ARL13B-dependent targets. To address this challenge, we incorporated both independently-derived mutant clones and matched rescue cell lines into our experimental design. Rescue analysis allowed us to filter transcriptional changes that were unrelated to loss of ciliary ARL13B function, while comparison across multiple mutant-rescue pairs enabled us to identify genes whose expression consistently tracked with ciliary ARL13B loss and restoration. This strategy reduced more than one thousand candidate genes identified in individual analyses to a core set of 131 overlapping DEGs, generating a high-confidence list of candidate downstream targets.

Several limitations should be considered when interpreting these findings. First, each rescue line represents an independently-derived clonal population, and we did not generate wildtype cells expressing the ARL13B-V5-His transgene or ARL13B-null cells to include in the RNA-seq analysis. Although the consistency of transcriptional changes across matched mutant-rescue pairs argues against clonal or transgene-related effects as the primary explanation for the identified DEGs, we cannot exclude the possibility of some contribution by these factors. Second, the rescue cell lines appear to reflect partial rather than complete restoration of ciliary ARL13B function, as demonstrated by incomplete downstream ciliary phenotypes, including ARL3 localization. It is not clear whether this is due to ARL13B-V5-His expression being driven by a non-endogenous promoter or something about the selection process of the cell lines. Regardless of the reason, it implies that the 131-gene set identified here may represent only a subset of ciliary ARL13B-dependent transcriptional responses. Consistent with this interpretation, many genes showed only partial normalization toward wildtype expression. While the 131 overlapping DEGs represent the highest-confidence, ciliary ARL13B-responsive candidates, partially rescued genes may also represent biologically relevant consequences of reduced ciliary ARL13B function and warrant future investigation.

Beyond the specific biological insights gained here, these findings underscore the value of combining independent CRISPR clones with genetic rescue approaches in exploratory transcriptomic studies. Prioritizing changes that are both reproducible across clones and reversible upon rescue strengthens causal inference and provides a practical framework for distinguishing biologically meaningful signals from clone-specific noise.

In conclusion, our study demonstrates that ciliary ARL13B is dispensable for ciliogenesis in kidney epithelial cells but is required to maintain normal ciliary composition and associated transcriptional programs. The resulting gene expression changes identify candidate pathways through which disruption of ciliary ARL13B may contribute to renal cystogenesis and further support the emerging concept that ciliary signaling defects, rather than loss of cilia themselves, can drive kidney disease (Piontek et al. 2007; Van Sciver et al. 2023; Pazour et al. 2002; Yoder et al. 2002). More broadly, our work highlights the importance of ciliary composition as a regulator of nuclear responses and establishes a methodological framework for improving confidence in RNA-seq analyses of CRISPR-derived cell models. These findings provide a foundation for future studies aimed at defining the signaling pathways that connect ciliary ARL13B to transcriptional regulation and kidney homeostasis.

## Data Availability

All processed expression data are available in supplementary tables. Bulk RNA sequencing data is available on GEO at GSE342252. All software used in this work is described in the Methods section and data made publicly available via Figshare at 10.6084/m9.figshare.c.8650158. Cell lines and plasmids are available upon request.

## Acknowledgements

This work was supported by a grant from the National Science Foundation (2329634). We are grateful to members of the Caspary lab as well as Quinn Eastman for feedback, comments, and suggestions in the preparation of this manuscript.

## Notes

https://doi.org/10.6084/m9.figshare.33278820

### Competing Interest Statement

The authors have declared no competing interest.

### Summary of Updates

Ciliary intensity and statistical analysis updates during manuscript revision process

