## Supplementary figures and images for "Transcriptomics of independent CRISPR-edited cell lines reveal ciliary-specific ARL13B dependent changes"

### Supplemental Figure 1

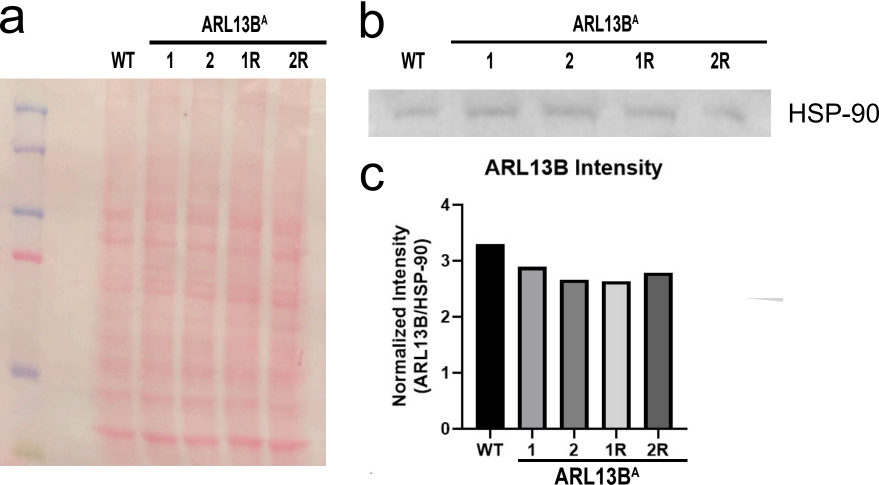

### Supplemental Figure 2

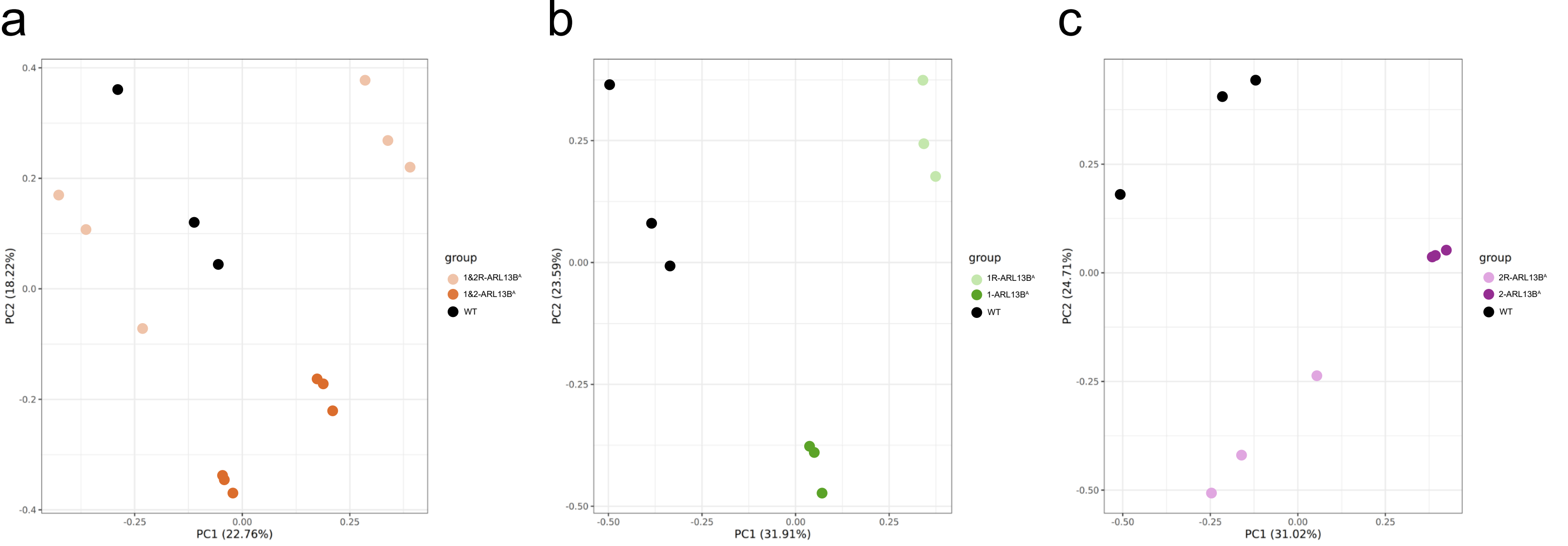

### Supplemental Figure 3

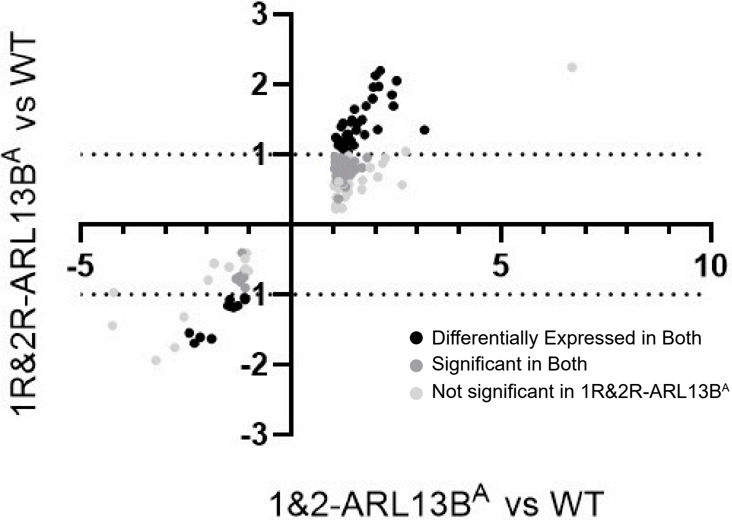
